# A time-resolved atlas of histone modifications during mitotic entry

**DOI:** 10.1101/2025.09.29.679131

**Authors:** Natalia Y. Kochanova, Marco Borsò, Moonmoon Deb, Shaun Webb, Ipek Ustun, Kumiko Samejima, Ignasi Forne, Itaru Samejima, Linfeng Xie, James R. Paulson, Axel Imhof, William C. Earnshaw

**Author notes:** Equal contribution (Natalia Y. Kochanova, Marco Borsò, Moonmoon Deb).

## Abstract

Mitotic chromosome formation is essential for faithful chromosome segregation in metazoans. While condensin complexes are critical for the formation of rod-shaped mitotic chromosomes, additional mechanisms—particularly those involving phosphorylation and deacetylation of specific histone residues—have been proposed to contribute a further 2- to 4-fold reduction in mitotic chromatin volume. In this study, we employ high-resolution mass spectrometry to determine the kinetics of histone modifications in cell cultures undergoing a highly synchronous mitotic entry at 2.5-minute resolution. Our analysis reveals three different programmes of histone H3 phosphorylation on T3, S10 and S28. These modifications are consistent with methyl-phos switches regulating the association of readers with chromatin other than at promoters. Mass spectrometry and quantitative ChIP-Seq reveal that H3 T3 phosphorylation is a general marker of heterochromatin and not specifically centromeres as previously suggested. Finally, we show that histone acetylation undergoes only modest changes as rod-shaped chromosomes form during unperturbed mitotic entry. Thus, previously reported reductions in acetylation associated with chromosome formation were apparently attributable to delays in mitotic exit used as part of mitotic synchronisation protocols. The mechanism of condensin-independent chromatin compaction in mitosis remains unexplained.

## Introduction

DNA in eukaryotic cells is packaged in nucleosomes containing an octamer of histone proteins with two molecules each of H2A, H2B, H3 and H4 (Luger et al., 1997). Access to the DNA is regulated by histone modifications deposited, erased and read by a wide variety of proteins and protein families (Kouzarides, 2007, Taverna et al., 2007, Hyun et al., 2017). Differing modifications mark different functional classes of chromatin (Filion et al., 2010, Encode Project Consortium et al., 2020, Millan-Zambrano et al., 2022). Histone modification patterns are regulated throughout the cell cycle (Bonenfant et al., 2007, Alabert et al., 2015, Lu et al., 2021) and are thought to change dramatically in mitosis (Garcia et al., 2005, Bonenfant et al., 2007, Lin et al., 2016, Zhiteneva et al., 2017, Javasky et al., 2018, Behera et al., 2019, Lu et al., 2021) as chromosomes condense and form highly compact rod-shaped mitotic chromosomes (Earnshaw, 1991, Swedlow and Hirano, 2003, Paulson et al., 2021, Hirano, 2025).

The best known modification of core histones in mitosis is phosphorylation. Phosphorylation of histone H1 was the classic biochemical assay used in the identification of Cdk1-cyclin B as the kinase that triggers mitotic entry (Lohka et al., 1988). (We do not discuss this modification further here as H1 is not present in our histone preparations). Phosphorylation of histone H3 Serine 10 (H3S10ph) during interphase is confined to active promoters in Drosophila (Ivaldi et al., 2007) and stem cells (Tiwari et al., 2011) at very low levels. H3S10 phosphorylation is much more prominent in mitosis (Allis and Gorovsky, 1981, Wei et al., 1998). This mark dramatically accumulates starting in prophase, peaks at metaphase and declines on ana/telophase chromosomes as revealed by fluorescence microscopy (Hendzel et al., 1997), western blotting (Qian et al., 2011) or mass spectrometry (Lin et al., 2016). These kinetics were confirmed in living cells by FabLEM (Hayashi-Takanaka et al., 2009, Stasevich et al., 2014, Ruppert et al., 2018). Kinetics of other histone phospho marks has also been studied in mitosis, albeit, as in the case of H3S10ph with limited cell synchrony (Dai et al., 2005, Qian et al., 2011, Lin et al., 2016).

H3S10ph is deposited by the chromosomal passenger complex (CPC) kinase Aurora B (Earnshaw and Bernat, 1991, Hsu et al., 2000, Adams et al., 2000), which also phosphorylates Serine 28 (Goto et al., 1999, Goto et al., 2002, Watson et al., 2020). Aurora A also phosphorylates H3S10ph (Crosio et al., 2002). Histone H3 is also phosphorylated on Threonine 3 (H3T3ph) by Haspin kinase (Dai et al., 2005, Watson et al., 2020). H3T3ph was reported to be necessary for CPC recruitment to the inner centromere (Kelly et al., 2010, Wang et al., 2010, Yamagishi et al., 2010, Jeyaprakash et al., 2011). In addition, the H3.3 isoform is phosphorylated on Serine 31 (Hake et al., 2005) by Chk1 (Chang et al., 2015, Day et al., 2025) and histone H2A is phosphorylated on Threonine 120 (H2AT120ph) by Bub1. This latter phosphorylation, together with H3T3ph, participates in the centromeric localization of Shugoshin and the CPC (Kawashima et al., 2010, Yamagishi et al., 2010).

Phosphorylation sites of T3, S10, and S28 on histone H3 are adjacent to the methylated lysines at K4, K9 and K27. Studies of histone methylation in mitosis have yielded contradictory results. For example, H3K9 methylation has variously been reported to increase (McManus et al., 2006), decrease (Peng et al., 2018) or remain unaltered during mitosis (Fischle et al., 2005). Importantly, those studies used fluorescent reporters or western blotting to track the levels of histone methylation, and it has been reported that S10 phosphorylation can alter antibody recognition of H3K9 methylation (Duan et al., 2008). Thus, it is important to use orthogonal methods, such as mass spectrometry to confirm the fate of phosphorylation and other modifications of key chromosomal proteins, including histones in mitosis (Ohta et al., 2016, Zhiteneva et al., 2017, Javasky et al., 2018).

Many writers, erasers and readers dissociate from chromosomes during mitosis, and it has been suggested that mitosis is effectively a time of “cleaning up” the genome (Yanagida, 2009). It was suggested that some methylated lysine readers (for example, HP1α) dissociate from their binding sites once T3, S10 and S28 are phosphorylated (the so-called methyl-phos switch hypothesis). This was first proposed for HP1 ejection from H3K9me3 following phosphorylation of H3S10 (Fischle et al., 2005, Hirota et al., 2005) (although see (Terada, 2006)), and subsequently reported for ING1, DIDO and PHF8 being displaced from H3K4me3 by phosphorylation of T3 (Gatchalian et al., 2016) and *Drosophila* PH and PC being displaced from H3K27me3 following phosphorylation of S28 (Fonseca et al., 2012). This hypothesis was challenged for Lys4 of H3 following the observation that Haspin knockout does not induce reassociation of a number of H3K4me3 readers with mitotic chromosomes together with the result that H3T3ph and H3K4me2/3 appear by ChIP-seq to be mutually exclusive on a genome-wide level (Harris et al., 2023). Given that H3T3 is not phosphorylated on peptides containing H3K4me2/3 ((Garcia et al., 2005) and confirmed by our quantitative analysis below), the methyl-phos hypothesis clearly could not apply to the readers of this residue in the di-and tri- methylated state, though it could apply for readers of H3K4me1.

Histone deacetylation has also been proposed to play a role in mitotic chromatin compaction (Zhiteneva et al., 2017, Gandhi et al., 2022). Acetylation levels of mitotic histones purified from nocodazole- or STLC-blocked cells are largely decreased (Zhiteneva et al., 2017, Javasky et al., 2018) or moderately decreased at some sites (Behera et al., 2019). *In vitro* acetylation reported to decompact chromatin (Hizume et al., 2010) by neutralizing positive charges on the histone N-termini and decreasing histone affinity for DNA (Nitsch et al., 2021). Although acute condensin depletion did not change the over-all chromatin volume in mitosis (Samejima et al., 2018), it was reported that treatment with histone deacetylase (HDAC) inhibitor trichostatin A (TSA) led to a dramatic decompaction of mitotic chromatin after simultaneous acute depletion of condensin (Schneider et al., 2022). Furthermore, nucleosome deacetylation also increased the propensity of isolated nucleosomes to undergo phase separation in vitro (Gibson et al., 2019). This and other experiments led to the proposal that histone deacetylation is a key factor in the compaction of mitotic chromosomes (Wilkins et al., 2014, Kruitwagen et al., 2015, Schneider et al., 2022). However, TSA influences the acetylation of multiple proteins, including cell cycle regulators such as p53 (Wood et al., 2018). Therefore, the question of whether deacetylation of histones plays a role in chromosome compaction merits further investigation.

Here, we have probed the kinetics of core histone phosphorylation, acetylation and methylation in vertebrate cells with minute-by-minute resolution while cells progress freely through prophase and prometaphase. To achieve synchronous free-running mitotic entry, we utilized a Cdk1^as^ chemical-genetic system in chicken DT40 cells (Hochegger et al., 2007, Gibcus et al., 2018, Samejima et al., 2018, Samejima et al., 2022, Samejima et al., 2025). We analysed the interplay of selected phospho- and methyl marks by liquid chromatography-mass spectrometry (LC-MS) and mapped the location of key marks on the DT40 genome using ChIP-sequencing. Overall, this allowed us to create a dynamic atlas of histone marks with unprecedented temporal resolution as mitotic chromosomes are formed. We observed 3 differing phosphorylation programs throughout mitotic entry and show that H3T3 phosphorylation is absent at transcription start sites. On a genome-wide level, our ChIP-seq findings indicate that H3T3ph is heterochromatic and is depleted at non-repetitive centromeres. Additionally, we find that core histones are only mildly deacetylated as rod-shaped chromosomes compact in unperturbed prophase, but that acetylation levels drop dramatically if mitotic exit is inhibited by nocodazole or MG132.

## Results

### Rapid collection of cells throughout mitotic entry for the purification of histones allows measurements of the kinetics of histone marks

For histone purification, we designed a small-scale variant of a previously published protocol, which allows full preservation of phosphorylation marks on purified core histones (Rodriguez-Collazo et al., 2009, Leidecker et al., 2016, Zhiteneva et al., 2017).

The chicken DT40 Cdk1^as^ cell line undergoes a highly synchronous mitotic entry within minutes of release from a G_2_ arrest with 1NM-PP1 (Hochegger et al., 2007, Gibcus et al., 2018, Samejima et al., 2018). After washout of 1NM-PP1, DT40 Cdk1^as^ cells enter mitosis within minutes. This allows us to obtain cell populations entering prophase and forming mitotic chromosomes with minute-by minute precision. This same technology can be applied to various human cell lines, but in general their entry into mitosis is delayed by tens of minutes. Therefore, the degree of mitotic synchrony (which is well over 90% in prophase, for example) means that DT40 cells offer a highly advantageous system to study the biochemistry of mitotic chromosome formation. Following 1NM-PP1 washout we collected samples at 0.5, 2.5, 5, 7.5, 15 and 30 minutes into mitosis (Fig. 1A). The nuclear lamina was still present at the 7.5-minute timepoint (Fig. 1B). We therefore classified the first 4 timepoints as prophase, and the 15- and 30-minute timepoints as prometaphase, based on Lamin B1 immunofluorescence staining (Fig. 1C).

**Fig. 1.**
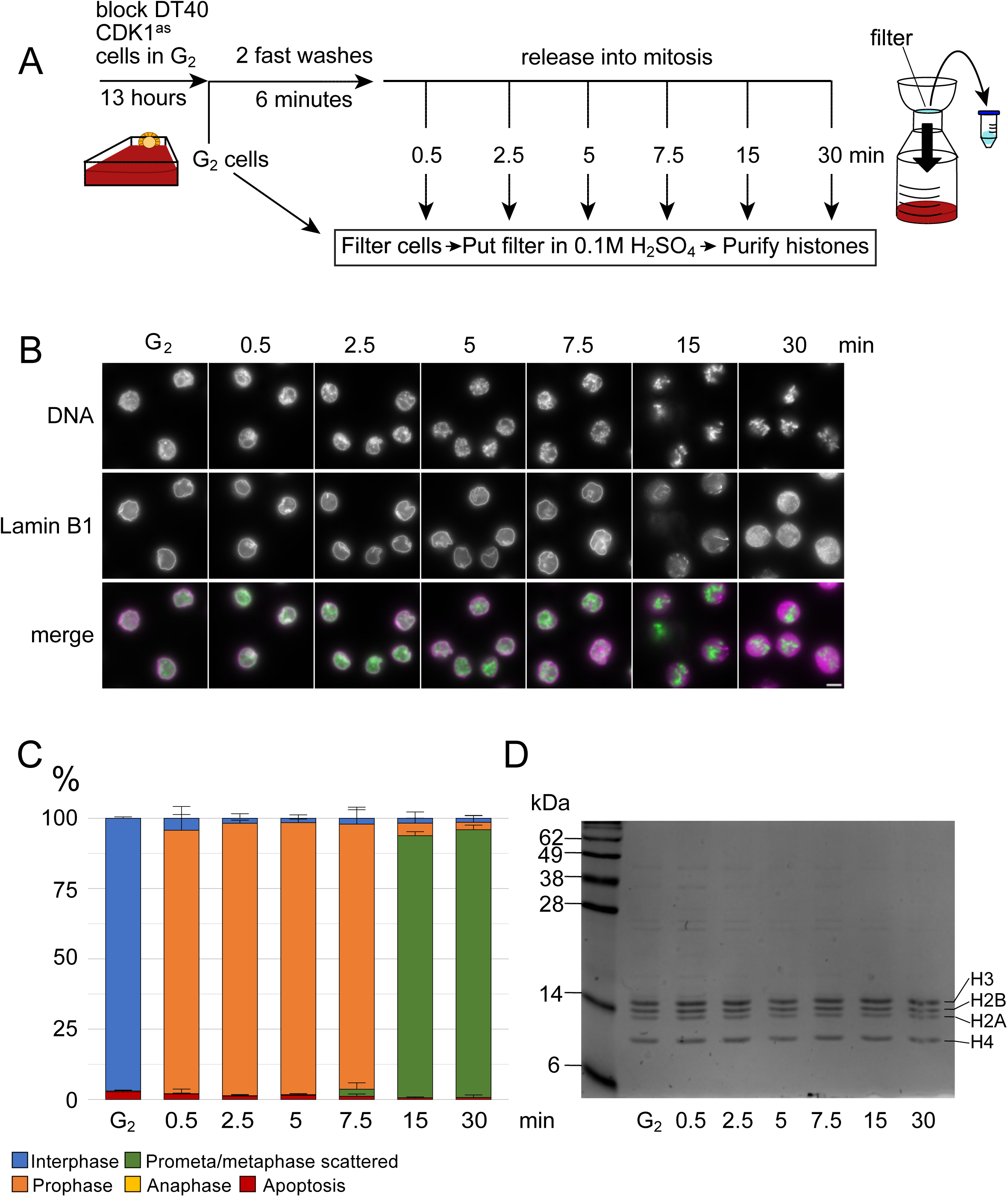
Purification of histones throughout synchronous mitotic entry. A – A scheme of the free-running mitotic entry experiment. B – Images from the experiment with immunofluorescence against Lamin B1. Scale bar represents 5 μm. C – Quantification of mitotic phases from B. Bars represent mean ± standard deviation. n=4. D – Coomassie stained gel of purified histone proteins from the experiment.

The cell suspension was harvested by filtration on a 0.2-μm nitrocellulose membrane during the last 30 seconds of each timepoint. This filter was stored in 0.1 M sulfuric acid on ice. Histones were isolated by acid extraction, purified by small-scale cation exchange chromatography, resolved on SDS-PAGE (Fig. 1D) and excised from the gel for mass spectrometry (Alabert et al., 2015, Volker-Albert et al., 2018).

To compare our results with published studies in which mitotic cells were accumulated using nocodazole or STLC block, we released Cdk1^as^ cells into early prometaphase and then added nocodazole for 2 hours after release, before harvesting cells and purifying histones as described above (Fig. S1A, B, C, D). In nocodazole, prometaphase cells displayed a “compact” phenotype with the chromosomes collapsed into a ball, in contrast to “scattered” prometaphase without nocodazole (Fig. S1B, C). This is likely due to clustering around centrosomes as kinetochores attach to short microtubule fragments (our unpublished results). This methodology allowed us to profile the kinetics of histone modifications with a minute-by-minute resolution during mitotic entry.

### Phosphorylation is the only histone modification to change dramatically during mitotic entry

Our system allowed us to map the kinetics of histone marks during mitotic entry to an extent not previously possible due to lack of high synchrony. We used Skyline (Pino et al., 2020) to quantify 60 core histone peptides bearing differing modifications throughout mitotic entry (Table S1). For each modification, we calculated a percentage by summing the intensities of all forms of a given peptide bearing the indicated modification divided by the sum total of all intensities of that peptide regardless of modification status (Fig. 2A).

**Fig. 2.**
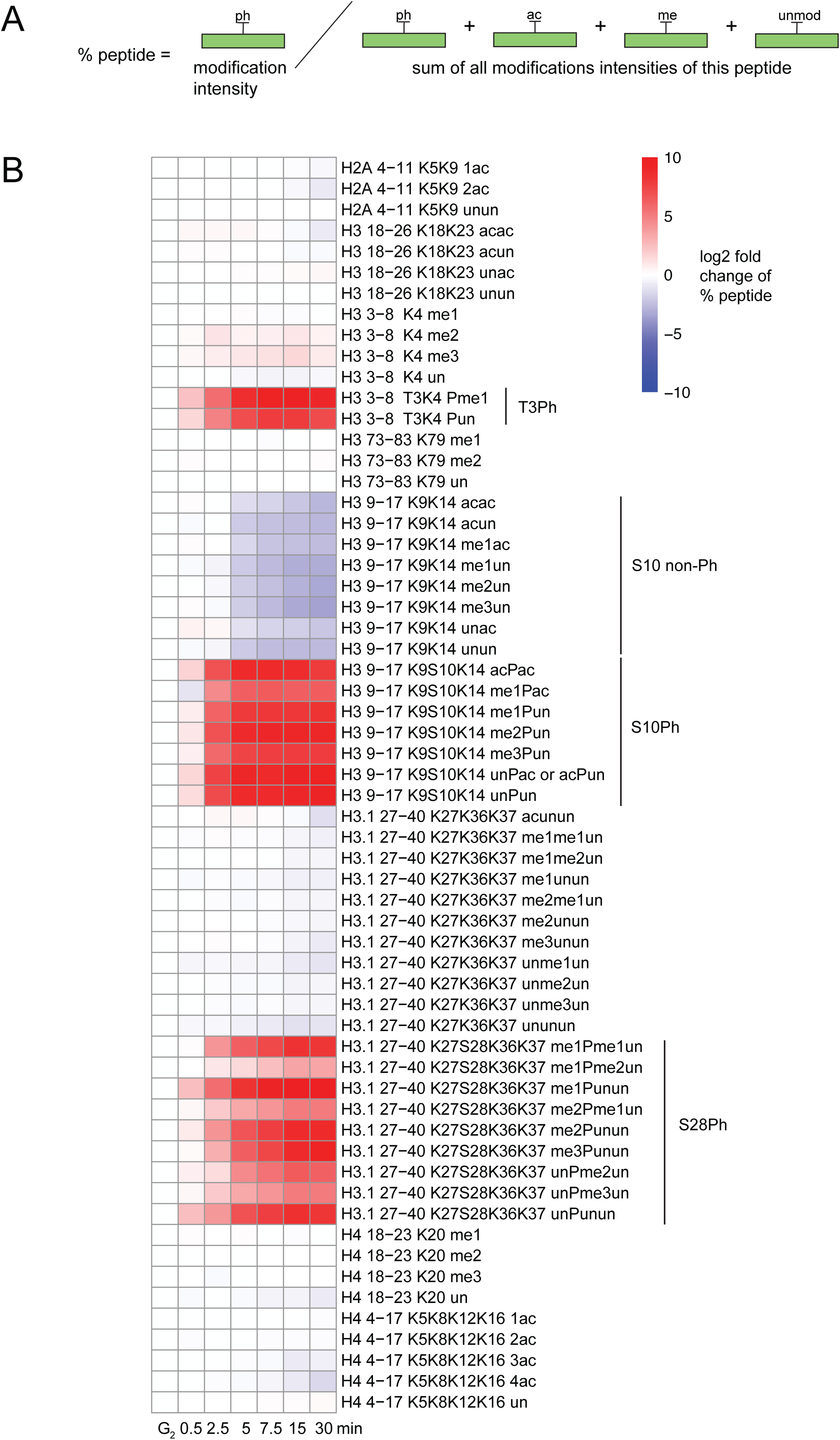
An atlas of changes in histone modifications throughout mitotic entry. A – A scheme of peptide % calculation. It is computed as the modification intensity divided by the sum of all intensities of differing modifications of this peptide. B – A heatmap representing log_2_fold change of mean peptide % relative to G_2_ timepoint during the free-running mitotic entry. 60 peptides quantified in this study are represented. N=4.

To display trends as cells entered mitosis, we plotted a heatmap displaying the mean log_2_fold change of histone modification levels relative to the G_2_ timepoint (Fig. 2B). This exhibited a striking rise in the levels of phosphorylated peptides and a decline in their non-phosphorylated counterparts. The levels of all other histone modifications were largely unchanged during mitotic chromosome compaction. As expected, interphase histone modification levels (G_2_ timepoint) were consistent with a previous study on human cells (Alabert et al., 2015) (Table S2).

### Three different programs of phosphorylation occur during mitotic entry

As expected from previous studies, phosphorylation on serine 10 of histone H3 gradually increased during prophase and further in prometaphase, reaching 74.2 ± 2.7 % of peptide by 5 minutes and 88.9 ± 0.5 % of peptide by 30 minutes (Fig. 3A). This high level of modification confirms the effectiveness of our isolation protocol in preserving phosphorylation of the core histones. In all cases, phosphorylation levels in nocodazole treated cells were generally comparable to the levels of late prometaphase (30-minute timepoint) in free-running mitosis (Fig. 3 and S2A-C, Table S3).

**Fig. 3.**
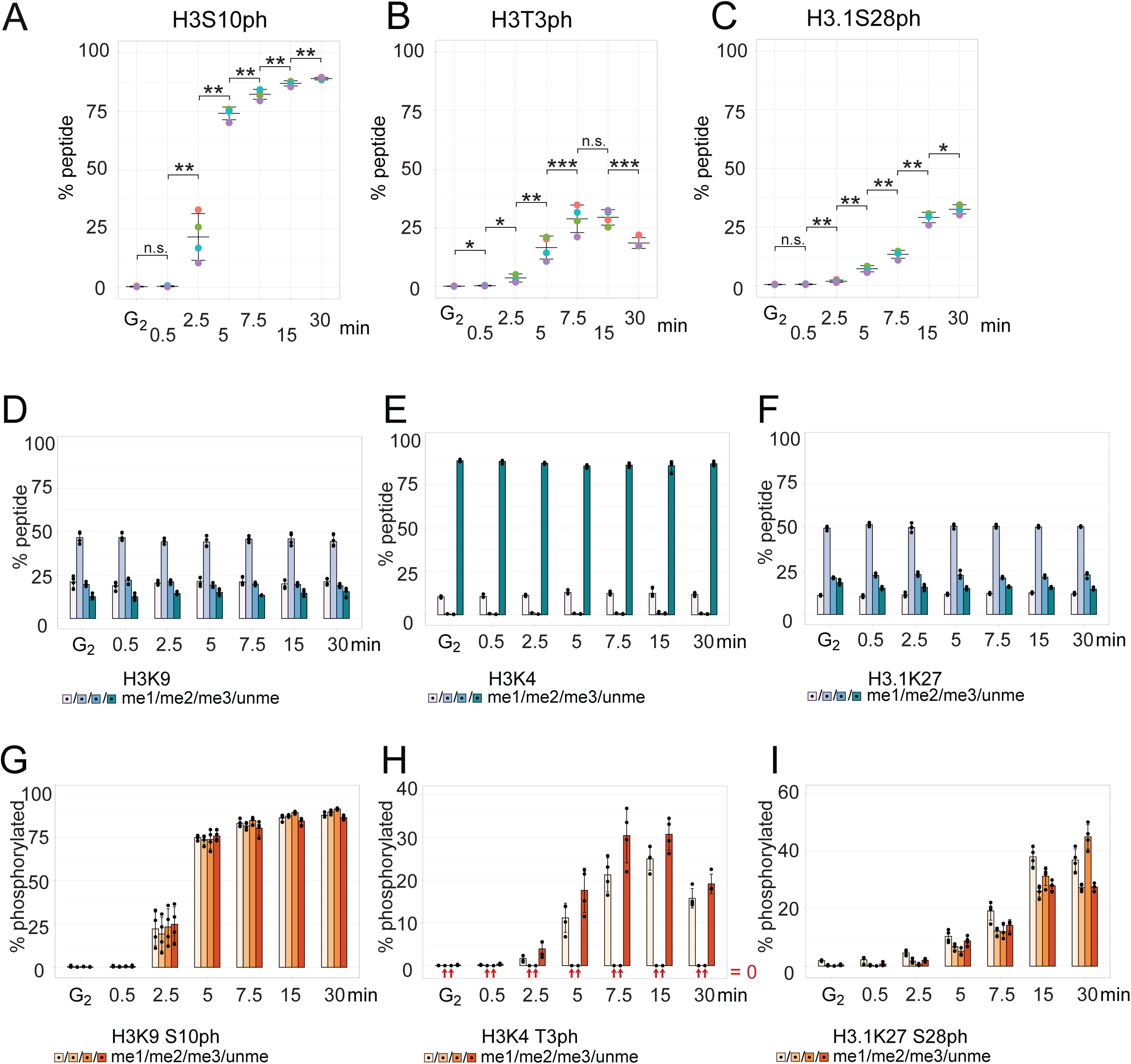
Three different programs of phosphorylation throughout mitotic entry. A, B and C – H3S10, H3T3 and H3.1S28 phosphorylation levels as the % of total peptide in the free-running mitotic entry experiment. Bars represent mean ± standard deviation. Individual replicates are shown with distinct colors. N=4. For statistical comparison, Kruskal-Wallis test, followed by Conover-Iman test (in case of p < 0.05 in Kruskal-Wallis test) was performed. The p-values were adjusted with Benjamini-Hochberg procedure. D – A grouped barplot of the total mean peptide % of H3K9me1, me2, me3 and unme modifications in the free-running mitotic entry experiment. In the case of unmethylated peptide, we also included acetylation on K9 and K27, although generally the levels of acetylation were much lower than methylation levels (Table S1, Table S3). Bars represent mean ± standard deviation. n=4. G – A grouped barplot of the % of H3K9me1, me2, me3 and unme which is phosphorylated (% of total phosphorylated peptides normalized to the total peptide % of these modifications). Bars represent mean ± standard deviation. n=4. E and H – The same as in D and G, but for H3K4me1, me2, me3 and unme. For H3K4me2 peptide % at 15 min an outlier replicate was excluded from the calculation of mean and standard deviation. F and I – The same as in D/G and E/H, but for H3.1K27me1, me2, me3 and unme.

In contrast, phosphorylation of threonine 3 on histone H3 was both delayed and transient – the modification level reached 16.6 ± 4.9 % of the peptide at 5 min, peaked at 15 minutes of mitotic entry at 29.4 ± 3.3 % of the peptide and declined at 30 minutes to 18.5 ± 2.3 % of the peptide (Fig. 3B). These results are consistent with results of a previous study in which we used FabLEM (Stasevich et al., 2014) to demonstrate that during free-running mitosis in unsynchronized HeLa cells, H3T3 phosphorylation levels rise significantly later than and then decrease significantly before H3S10 phosphorylation (Ruppert et al., 2018).

Phosphorylation of serine 28 on H3.1 shows a third temporal pattern. This phosphorylation is delayed in early prophase in comparison to the other two phospho-marks, reaching 7.2 ± 1.3 % of the peptide at 5 min. It then steadily increases to 32.5 ± 2.0 % of peptide at 30 minutes (Fig. 3C). Interestingly, if cells are delayed in mitosis with nocodazole or MG132, H3S28ph never rises above the level of ∼30% of total peptide. Similar to T3ph, H3S28ph shows a decline in cells held in mitosis for 120 minutes with nocodazole (Fig. S2B and C, Tables S3 and S4). This is in contrast to S10 phosphorylation, which rises rapidly to near quantitative levels and then remains high throughout a 120-minute mitotic arrest (Fig. S2A).

The high enrichment of phospho marks throughout mitosis observed in our study agrees with one previous study in which 80% of H3S10 peptide was phosphorylated in cells arrested in prometaphase with STLC (Javasky et al., 2018), but is in disagreement with two previous LC-MS reports in which H3S10ph levels reached maximum peptide levels of approx. 50%, and H3T3ph and H3S28ph were in the maximum range of approx. 2-8% (Lin et al., 2016, Lu et al., 2021). Disagreements with previous results might reflect losses of phosphorylation during histone isolation, synchronization- and/or species-specific differences or technical differences between bottom-up and middle-down MS analysis. In general, however, it is difficult to imagine how technical issues would increase, rather than decrease phosphorylation efficiency.

We conclude that the three phosphorylation events on Histone H3 assayed here show distinct programs in early mitosis.

### Phosphorylation marks exhibit differing associations with methylated peptides

Strikingly, phosphorylated residues S10, T3 and S28 are adjacent to methylated residues K9, K4 and K27, all of which are bound by protein readers that influence chromatin activity and are inactive during mitosis. This led to the methyl-phos switch hypothesis, proposing that phosphorylation evicts methylation readers from adjacent residues (Fischle et al., 2005, Hirota et al., 2005, Fonseca et al., 2012, Gatchalian et al., 2016). Our LC-MS approach allowed us to measure adjacent phosphorylation and methylation events on the same peptide, thereby allowing us to determine the interplay of these marks.

Levels of both constitutive and facultative heterochromatin do not change during mitotic entry, and indeed, our data suggest that they have no effect on the program of mitotic phosphorylation of histone H3. We observed no major change in the levels of H3K9me1, me2, me3 and unme peptides (or H3.1K27me1, me2, me3 and unme peptides) during mitotic entry (Figs. 2B, 3D and F). These observations are consistent with a study in HeLa S3 cells using spike-in peptides (Javasky et al., 2018) and a study on mouse erythroid cells (Behera et al., 2019), however another study using a different LC-MS approach reported an increase in H3K9me3 in mitosis (Lu et al., 2021).

All four classes of peptide were phosphorylated with comparable efficiencies on both S10 and S28 (Fig. 3 and S2 G,I). Bearing in mind that S10 phosphorylation is at all time points much more extensive than S28 phosphorylation the overall levels were different but, importantly, there was no selectivity of these phosphorylation events occurring on either methylated or unmethylated K9/K27. Thus, both S10 and S28 phosphorylation essentially “paint” the entire genome, although the extent of “painting” by S10ph is significantly greater. The reason for the difference in levels and timing of the two phosphorylation events is not known.

The levels of H3K4me1, me2, me3 and unme also did not exhibit major changes throughout mitotic entry (Fig. 3E, Fig. S3A, C), consistent with a report on STLC-blocked cells (Javasky et al., 2018) but contrary to a previous study on mouse erythroid cells which reported an increase of H3K4me3 levels in mitosis (Behera et al., 2019). H3K4me2 and me3 containing peptides constituted a very small percentage of the overall peptide (0.55 ± 0.04 % for me2 and 0.25 ± 0.07 % for me3 in G_2_). This might seem surprising, given the many ChIP-sequencing studies describing these marks as being present at promoters genome-wide (Encode Project Consortium, 2012). However, promoters comprise only a small fraction of the genome. Indeed, the low levels of these marks measured by LC-MS in this study agree with previous mass spectrometry results from *Drosophila* embryos (Bonnet et al., 2019) and with a previous report on HeLa S3 cells (Javasky et al., 2018). Overall, we only detect H3K4 methylation on approximately 10% of histones (Fig. 3E).

We failed to detect any phosphorylation of H3K4me2- and me3-containing histones during mitotic entry in agreement with previous mass spectrometry (Garcia et al., 2005) and ChIP-Seq (Harris et al., 2023) studies (Fig. 3H). Phosphorylation was restricted to H3K4me1 and unmodified histones (see also (Garcia et al., 2005)), albeit less efficiently than H3K9 and H3.1K27 peptides (both methylated and unmethylated - Fig. S2G, H, I, Fig. S3B, D). As with overall levels of phospho marks, nocodazole-blocked cells in our study displayed the same phosphorylation/methylation trends, as the 30-minute timepoint of free-running mitotic entry (Fig. 3D-I, Fig. S2D-I).

In context of methyl-phospho switch hypothesis, some methylation readers have been shown to dissociate from mitotic chromosomes (Fischle et al., 2005, Hirota et al., 2005, Fonseca et al., 2012, Gatchalian et al., 2016). However, there has been no systematic description of the association/dissociation profiles of methyl readers on chromosomes in mitosis to date. To better understand the interplay between methyl/phospho marks and the relevant modifying enzymes and readers, we re-analysed the data set from (Samejima et al., 2022) to observe the dynamics of chromatin association during mitotic entry for H3K9me, H3K4me and H3K27me writers, erasers and readers (Binda and Fernandez-Zapico, 2016, Hyun et al., 2017, Jain et al., 2020, Sterling et al., 2021) (Fig. S4A-I). We observed that most these proteins dissociated from chromatin as cells entered mitosis, specifically at 10-15 minutes of mitotic entry (i.e. during or just after nuclear envelope breakdown). In contrast, Aurora B kinase, responsible for H3S10ph and H3S28ph deposition, did not dissociate from chromosomes (Fig. S4J). Haspin kinase (GSG2 in chicken) displayed a slight decrease in chromatin association during 10-15 minutes of mitotic entry in the dataset (Fig. S4J). Levels of PP1γ phosphatase, which dephosphorylates H3 residues (Qian et al., 2011), also exhibited a slight decrease on chromatin, while levels of its regulatory subunit, Repo-Man, fell more abruptly. Different subunits and regulators of PP2A, which is also involved in H3 dephosphorylation (Nowak et al., 2003), displayed differing kinetics of chromosome association throughout mitotic entry. Together, this analysis indicates that most histone H3K9, K4 and K27 methylation writers, erasers and readers dissociate from condensing mitotic chromosomes relative to their G_2_ abundance on chromatin, whereas most histone phospho- modifiers remain on mitotic chromosomes. This is consistent with the detected changes in H3T3, S10 and S28 phosphorylation and lack of changes in H3K4, K9 and K27 methylation during mitotic entry.

Taken together, we observed that the total levels of H3K4, H3K9, and H3.1K27 methylated and unmethylated peptides do not change during the entry into mitosis, but these peptides become efficiently phosphorylated on adjacent serine or threonine residues to differing extents. S10 and S28 phosphorylation are unaffected by methylation of the adjacent lysine, but T3 is never phosphorylated if Lys4 is di- or tri-methylated (see also (Garcia et al., 2005)). Since many readers recognize H3K4me2 and H3K4me3, this means that those readers are not affected by a phospho-methyl switch. This explains why Haspin knockout does not induce reassociation of a number of H3K4me3 readers with mitotic chromosomes (Harris et al., 2023).

### H3T3 phosphorylation avoids promoter-proximal active chromatin during mitotic entry

Our LC-MS approach allowed us to measure total levels of histone marks, however marks such as H3K4me2 and me3 are not abundant at the LC-MS level. This makes their precise quantification difficult due to a lower signal-to-noise level. To complement our LC-MS measurements, we therefore investigated the interplay between histone phosphorylation and methylation during mitotic entry on a genome-wide level, performing ChIP-sequencing for H3K4me1, H3K4me2, and H3K4me3 alongside ChIP-sequencing for H3T3 phosphorylation (± spike-in-normalization) (Fig. S5A, B and C). These datasets provide a genome-wide view of epigenomic landscapes throughout mitotic entry.

H3T3ph levels measured by ChIP-seq increased sharply within five minutes of mitotic entry, declined modestly at 7.5 minutes and 15 minutes, and rose again by 30 minutes (Fig. 4B, C). These temporal dynamics are consistent with our mass spectrometry analyses showing mitotic enrichment of H3T3ph (Fig. 2B and 3B). Pearson correlation analysis confirmed that H3T3ph was mutually exclusive with H3K4me2 and H3K4me3. In contrast, some colocalization was seen for H3K4me1 and H3T3ph (Fig. 4A). ChIP-seq profiles and enrichment peak counts for H3K4me1/2/3 remained largely stable during early mitosis (Fig. 4B, C), consistent with previous reports for HeLa, U2OS and RPE cells arrested in mitosis with STLC or nocodazole (Javasky et al., 2018, Kang et al., 2020, Harris et al., 2023). Minor discrepancies between H3T3ph LC-MS and ChIP-seq trends may reflect changes in epitope accessibility due to chromatin condensation or the fact that LC-MS addresses total histone levels while ChIP-seq measures enrichment on chromatin.

**Fig. 4.**
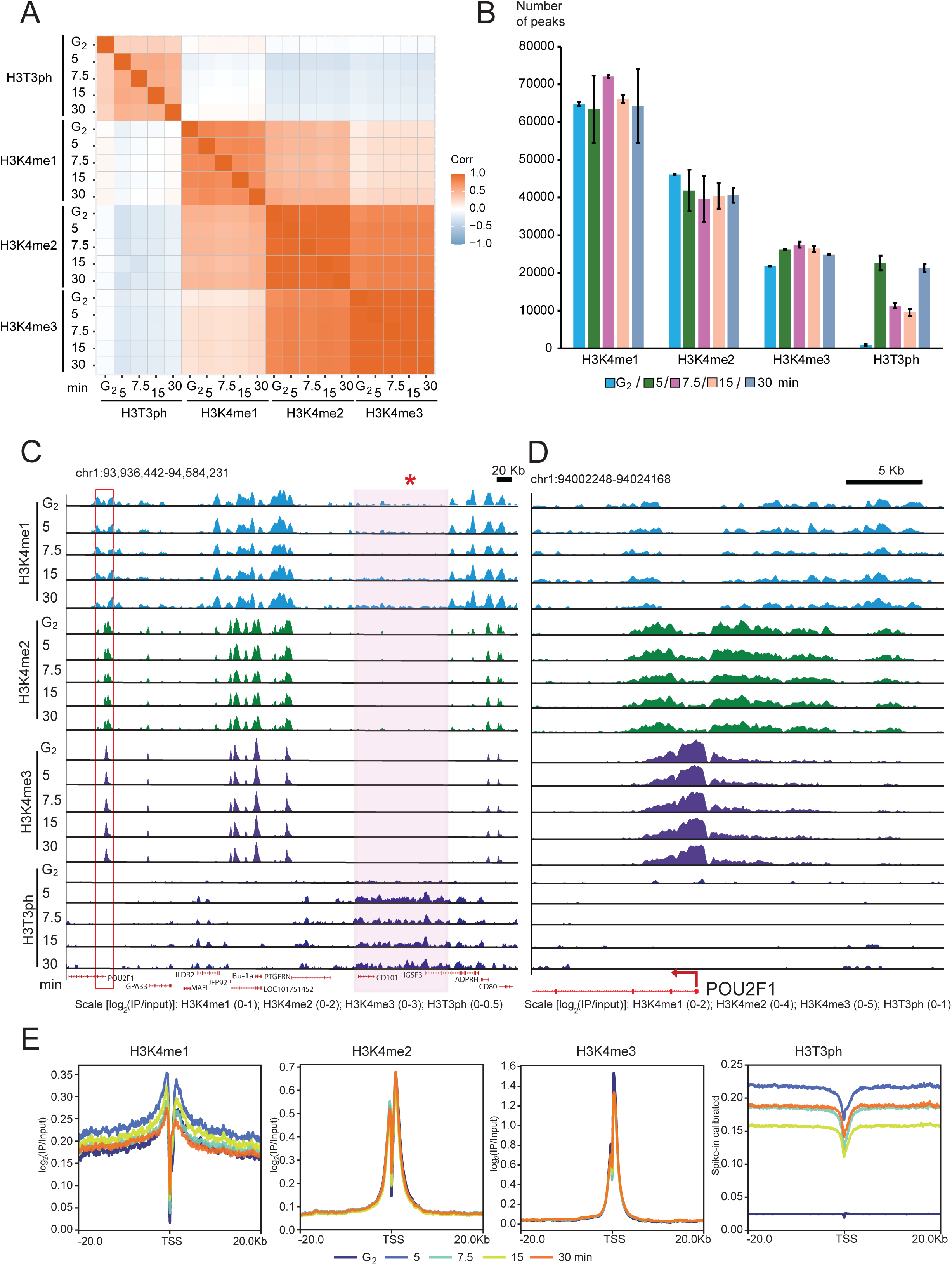
H3T3Ph is depleted from promoter proximal active chromatin during early mitosis. A – A Pearson correlation heatmap of ChIP-sequencing of H3T3ph, H3K4me1, H3K4me2 and H3K4me3 throughout mitotic entry. B – Numbers of enriched peaks at different timepoints. Bars represent mean ± standard deviation. n=2. C – A UCSC genome browser view showing the normalized (log_2_(IP/Input)) ChIP-seq read densities of H3K4me1 (light blue), H3K4me2 (dark green), H3K4me3 (violet) and H3T3ph (dark blue) on the chromosome 1. Regions shaded in pink (red *) highlight the overlap between H3K4me1 and H3T3ph. D – A magnified view of the red rectangle region from panel C, the TSS of *POU2F1* gene. E – Profile plots showing the normalized ChIP-seq signal densities of H3K4me1, H3K4me2, H3K4me3, and H3T3ph around the TSS (TSS ± 20 kb) of protein-coding genes.

Locus-specific analysis of a region near a transcription start site (TSS) on chromosome 1 confirmed well-known previous observations of H3K4me1/2/3 enrichment at promoter-proximal regions (Pokholok et al., 2005; Liu et al., 2005; Barski et al., 2007; Wang et al., 2008). H3K4me3 and H3K4me2 were concentrated at promoters, while H3K4me1 showed a broader distribution flanking promoter regions (Fig. 4D). Minor differences in the pattern of H3K4me2/3 around the TSS region (Fig. 4E) between this and previous reports (Javasky et al., 2018, Kang et al., 2020, Harris et al., 2023) might reflect the effect of nocodazole block or cell type-specific differences.

Notably, H3T3ph was absent from promoter-proximal regions. Correlation and enrichment analysis of our ChIP-Seq data confirmed the mutual exclusivity between H3T3ph and H3K4me2/3, consistent with our mass spectrometry analysis (Fig. 3H, Fig. 4A, C, D, E, Fig. S6, Fig. S8A) and with a prior ChIP-Seq study (Harris et al., 2023). Interestingly, unlike H3K4me2 and H3K4me3, we occasionally observed overlap between H3K4me1 and H3T3ph-enriched regions. This was most prominent at the 30-minute release time point (late prometaphase). However, the H3K4me1 signal at these overlapping sites was noticeably lower compared to regions with strong H3K4me1-enriched sites (Fig. 4C). Together, the ChIP data are in agreement with LC-MS results showing some T3 phosphorylation on H3K4me1 peptides, but not on H3K4me2/3, and with a previous report that Haspin activity is inhibited by H3K4 methylation (Eswaran et al., 2009). Thus, methyl/phos switching cannot apply to readers of H3K4me2/me3, but could apply for readers of H3K4me1.

Given the exclusion of H3T3ph from promoter-proximal active chromatin, we next investigated whether this modification preferentially associates with heterochromatin. To this end, we analysed the Pearson correlation of H3T3ph with well-established heterochromatic marks, H3K9me3 and H3K27me3 (Encode Project Consortium, 2012). Our analysis revealed a high correlation between H3T3ph and H3K9me3, a canonical mark of constitutive heterochromatin (Padeken et al., 2022). Using an unpublished dataset available on GEO (GSE242250), we detected no significant association of H3T3ph with H3K27me3 in DT40 cells (a marker for facultative heterochromatin) (Fig. 5A).

**Fig. 5.**
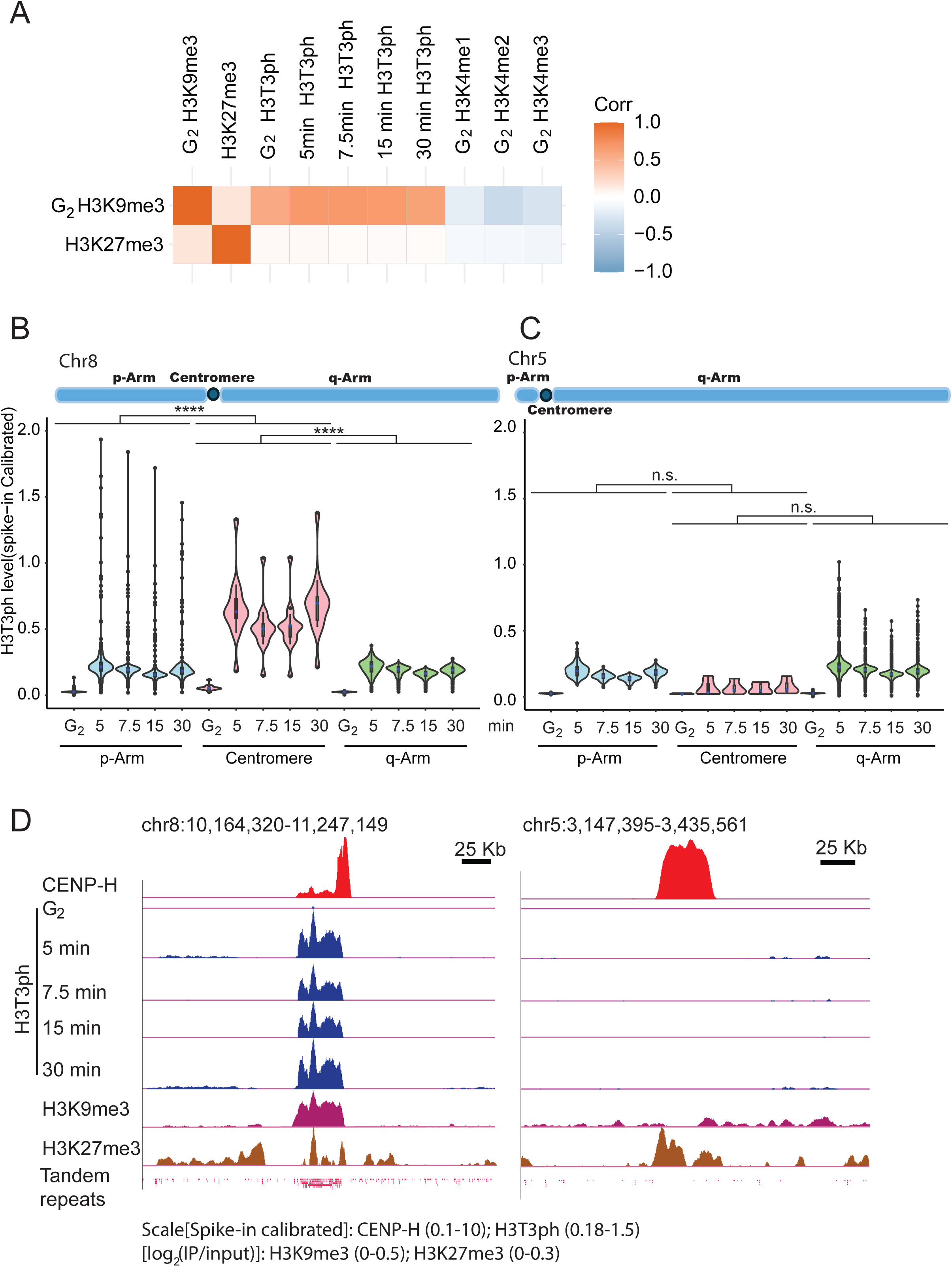
H3T3Ph distinguishes repetitive from non-repetitive centromeres. A – A Pearson correlation heatmap of ChIP-seq of H3T3ph throughout mitotic entry with H3K9me3 and H3K27me3. B – A violin plot of spike-in calibrated H3T3ph levels at the repetitive centromere of Chr8, as well as its p- and q-arms. C – The same but for Chr5 with the non-repetitive centromere. Statistics for B and C was computed by fitting a linear mixed effect model of log(enrichment). “Timepoint” and “position along the chromosome“ were fit as fixed effects, and “sample” as a random effect. Gaussian distribution of models’ residuals was confirmed. Since T-distribution is symmetric, one-sided p value was calculated for the null hypothesis of no centromeric enrichment of H3T3ph. The hypothesis was rejected in the case of a repetitive centromere on Chr8 (p < 0.0001, ****). The p-values were adjusted with Benjamini-Hochberg procedure. D – A UCSC genome browser view of the magnified centromeric region of Chr8 and Chr5. CENP-H ChIP-seq shows CENP-H (red)-binding peaks (spike-in calibrated) at the centromere of Chr8 (Left) and Chr5 (Right). Other tracks show landscape profile of H3T3ph (dark blue), H3K9me3 (dark pink) and H3K27me3 (dark orange) signal (spike-in calibrated or log_2_(IP/Input)) around the centromeric region.

Collectively, our findings indicate that H3T3ph undergoes global enrichment in heterochromatin during mitosis, while being selectively excluded from promoter regions characterised by active histone modifications such as H3K4me2/3.

### Functional centromeres without H3T3 phosphorylation

H3T3ph has previously been reported to be enriched at centromeres during mitosis, based on both indirect immunofluorescence (Dai et al., 2005, Kelly et al., 2010, Yamagishi et al., 2010, Harris et al., 2023) and ChIP-sequencing (Harris et al., 2023). This thought to be critical for localisation of the chromosomal passenger complex (Kelly et al., 2010, Wang et al., 2010, Yamagishi et al., 2010, Jeyaprakash et al., 2011). Centromeres are typically associated with heterochromatin, so it is difficult to distinguish between a role for T3ph in heterochromatin and at centromeres. The chicken genome presents a powerful model to distinguish between these two possibilities, as it contains three well-characterised non-repetitive centromeres on chromosomes 5, Z and W (Shang et al., 2010, Huang et al., 2023).

ChIP-seq profiling revealed robust enrichment of H3T3ph at repetitive centromeres during early mitosis, consistent with previous findings and its preference for heterochromatin (Fig. 5B and D, left). Unexpectedly, non-repetitive centromeres displayed no detectable H3T3ph signal (Fig. 5C and D, right; Fig. S7A and B). This was the case both in our original experiment and in a separate follow-up quantitative analysis using spike-in ChIP-seq (Fig. S5C). These findings confirm that H3T3ph is not universally required for centromeric identity or function (e.g. targeting of the CPC) (Kelly et al., 2010, Wang et al., 2010, Yamagishi et al., 2010, Jeyaprakash et al., 2011).

Genome-wide analysis revealed widespread enrichment of H3T3ph across the p- and q-arms of chromosomes during mitosis (Fig. S7A, S8A), indicating that its localization is not restricted to centromeric regions. Consistent with these findings, immunofluorescence imaging also demonstrated a broad chromosomal distribution of H3T3ph during mitotic entry (Fig. S8B).

Collectively, these findings demonstrate that non-repetitive centromeres can maintain centromeric function in the absence of H3T3ph despite the fact that repetitive centromeres consistently exhibit this modification. Furthermore, the widespread presence of H3T3ph across chromosomal arms suggests a broader mitotic role for this histone phosphorylation event than previously recognized.

### Histone acetylation changes little during mitotic entry as chromosomes form

Published experiments have led to a hypothesis that chromatin is deacetylated during mitotic entry, and that this drives or contributes to the compaction of mitotic chromatin (Wilkins et al., 2014, Kruitwagen et al., 2015, Zhiteneva et al., 2017, Javasky et al., 2018, Schneider et al., 2022). To understand the dynamics of histone acetylation as cells enter free-running mitosis in comparison to nocodazole block, we measured levels of selected acetylated peptides corresponding to each core histone under both conditions. We observed that monoacetylation on H2AK5/K9 and H3K9/K14, as well as diacetylation on H4K5/K8/K12/K16, were all either mildly decreased or unchanged during mitotic entry, but were profoundly reduced in cells released into mitosis and blocked with nocodazole for 2 hours (Fig. 6A, B and C). To generalize our findings, we plotted a heatmap of all quantified acetylated peptides and sums of peptides containing their phosphorylated and non-phosphorylated versions during time course and nocodazole experiments (Fig. 6D). While a few peptides such as H2AK5K9 diacetylation and H4K5K8K12K16 tetra acetylation displayed moderate decreases towards 30 minutes (a maximum -1.48 log_2_fold change during free-running mitosis), overall histone deacetylation was much more pronounced in nocodazole blocked cells with a maximum -4.37 log_2_fold change. Importantly, rod-shaped mitotic chromosomes were clearly seen at 5 and 7.5 minutes when acetylation levels were largely unchanged (Fig. 1B).

**Fig. 6.**
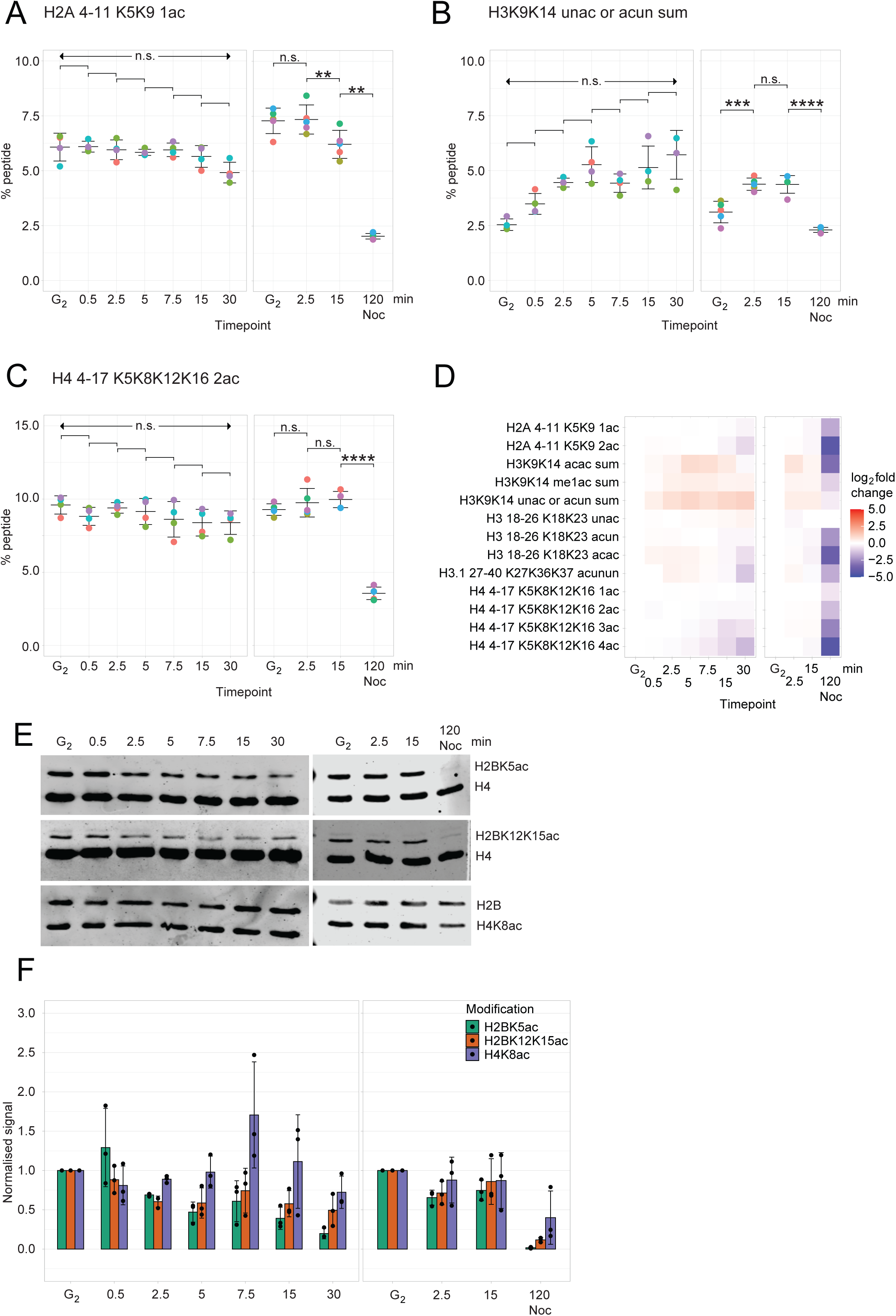
Acetylation is only minorly changed during chromosome compaction but declines in nocodazole treatment. A, B and C – H2A K5K9 1ac, H3 K9K14 unac or acun and H4 K5K8K12K16 2ac levels as the % of total peptide in the free-running mitotic entry (left) and nocodazole (right) experiments. Bars represent mean ± standard deviation. Individual replicates are shown with distinct colors. n=4 for the free-running mitotic entry experiment and n=5 for the nocodazole experiment. For statistical comparison, Kruskal-Wallis test, followed by Conover-Iman test (in case of p < 0.05 in Kruskal-Wallis test) was performed. The p-values were adjusted with Benjamini-Hochberg procedure. In case of a n.s. p-value in Kruskal-Wallis test or all n.s. p-values in Conover-Iman test, “n.s.” is shown for all adjacent timepoint comparisons. D – A heatmap representing mean log_2_fold change of acetylation levels of histones H2A, H3 and H4, as relative to G_2_. The data is represented for free-running mitotic entry and nocodazole experiments. Peptides denoted as “sum” were not measured as a whole, but represent the sum of measured % of all their ph and non-ph versions. E – Western blots against H2BK5ac/H4, H2BK12K15ac/H4 and H2B/H4K8ac for free-running mitotic entry (n=3) and nocodazole (n=3) experiments. H4 and H2B are used as loading controls. F – Quantification of E, normalized to G_2_ and loading control for each antibody and experiment.

It was technically not possible to detect H2B acetylation using our mass spectrometry pipeline. We therefore performed western blotting of the purified histones using antibodies recognizing H2BK5ac and K12K15ac in both free-running mitosis and nocodazole arrest experiments. The dynamics for both acetylation marks confirmed the trends observed for H2A, H3 and H4 by mass spectrometry. During free-running mitotic entry, these marks showed only a mild decrease, but their levels fell precipitously during the nocodazole block. Our LC-MS observations were also confirmed by western blotting of H4K8ac, although H2BK5ac and H2BK12K15ac displayed a more profound drop in nocodazole-treated samples compared to free-running mitotic entry (Fig. 6E, F).

Thus, in DT40 cells, histones are only mildly deacetylated as mitotic chromosomes are formed, but they are profoundly deacetylated when cells are delayed in mitosis by nocodazole treatment.

### Histones are gradually deacetylated as cells are delayed in mitosis

Our results suggest that extensive deacetylation of histones may be potentially caused by perturbing the microtubule cytoskeleton with nocodazole (as with the Eg5 inhibitor STLC (Javasky et al., 2018)) or a prolongation of mitosis as a consequence of checkpoint activation. To distinguish between these two possibilities, we released the cells in media containing either nocodazole or the APC/C inhibitor MG132 for 15, 30, 60 and 120 minutes, and measured histone modifications by LC-MS (Fig. S9A, B, C, and D, Table S4). Nocodazole treatment notably slowed mitotic entry with only a subset of cells having visibly disrupted nuclear envelope at 15 minutes, however the vast majority of cells progressed to prometaphase by 30 minutes and acquired the “compact” chromosome mass phenotype at 60 minutes (Fig. S9A B, C). MG132 - treated cells progressed into mitosis faster, and reached prometaphase already by 15 minutes, similar to the mitotic entry without perturbations (Fig. S9A B, C).

Mass spectrometry analysis revealed a progressive deacetylation of histones as cells are delayed in mitosis. The levels of acetylated peptides of histones H2A, H3 and H4 fell steadily during the delay (Fig. 7A-D). Thus, prolonged mitosis resulted in massive histone deacetylation in our system.

**Fig. 7.**
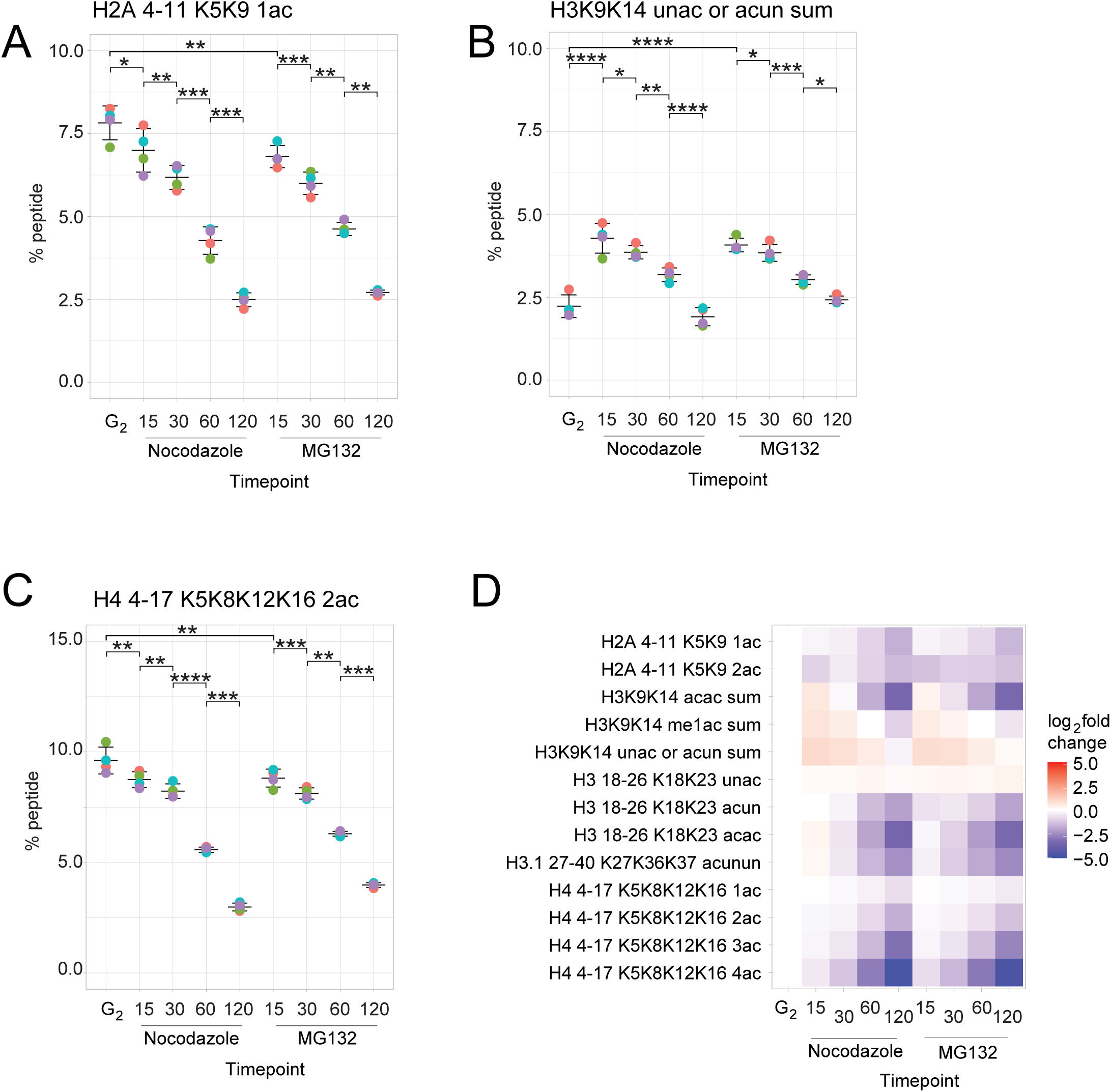
Acetylation gradually declines with increased length of mitosis upon checkpoints’ perturbation. A, B and C – H2A K5K9 1ac, H3 K9K14 unac or acun and H4 K5K8K12K16 2ac levels as the % of total peptide in the mitotic entry with nocodazole or MG132. Bars represent mean ± standard deviation. Individual replicates are shown with distinct colors. n=4. For statistical comparison, Kruskal-Wallis test, followed by Conover-Iman test (in case of p < 0.05 in Kruskal-Wallis test) was performed. The p-values were adjusted with Benjamini-Hochberg procedure. D - A heatmap representing mean log_2_fold change of acetylation levels of histones H2A, H3 and H4, as relative to G_2_. The data is represented for mitotic entry with nocodazole or MG132. Peptides denoted as “sum” were not measured as a whole, but represent the sum of measured % of all their ph and non-ph versions.

Taken together, our results suggest that previous measurements of histone modifications relying on collecting cells after mitotic arrest are likely to have led to overinterpretation of the magnitude of core histone deacetylation during mitotic chromosome formation.

## Discussion

Here we have used a chemical-genetic strategy (Hochegger et al., 2007, Gibcus et al., 2018, Samejima et al., 2018) to obtain populations of chicken DT40 cells entering mitosis with near-perfect synchrony. After washout of 1NM-PP1, DT40 Cdk1^as^ cells enter mitosis within minutes. This allowed us to to profile the kinetics of levels of 60 peptides on histones H2A, H3 and H4 throughout mitotic entry and upon mitotic delay.

This study has yielded several results that we will explore further here.

1. Histone methylation does not change as mitotic chromosomes are formed.
2. Histone phosphorylation changes dramatically, but significant differences are seen in both the kinetics and substrate specificity for different phosphorylation events.
3. Phosphorylation of H3T3 by haspin kinase does not specifically mark centromeres but instead is preferentially associated with heterochromatin.
4. Histone deacetylation occurs to only a very slight extent, if at all, during free-running mitotic chromosome formation. However, if cells are delayed in mitosis, histone acetylation falls precipitously.

Methylation marks associated with both active and inactive chromatin exhibit no significant changes during mitotic entry. Thus, the widely documented general inhibition of transcription that occurs at mitosis (Taylor, 1960, Parsons and Spencer, 1997, Palozola et al., 2017, Contreras and Perea-Resa, 2024) is not due to a loss of binding sites for chromatin readers in mitosis. Indeed, we confirm that writers, erasers and readers all dissociate from chromatin as chromosomes form in mid-late prophase. Importantly, readers could be released from chromatin during mitotic entry because they are themselves phosphorylated and have a lower affinity for chromatin. Alternatively, the phosphorylation of H3S10, which appears to be pervasive, effecting nearly 100% H3 molecules, could make chromatin generally more likely to eject bound factors.

An alternative and popular explanation is the methyl-phos switch hypothesis, originally proposed to regulate HP1 binding to H3K9me3 (Fischle et al., 2005, Hirota et al., 2005). This proposed that phosphorylation of S or T residues could displace readers bound to adjacent methylated K residues. However, this hypothesis was recently questioned for H3K4me2/me3 (Harris et al., 2023). Indeed we confirm that peptides containing di- or tri- methylated H3K4 are entirely devoid of phosphorylation during mitotic entry. H3K4me2/3 have been observed at promoters since the first ChIP-based experiments (Pokholok et al., 2005, Liu et al., 2005, Barski et al., 2007, Wang et al., 2008), and H3K4me3 was recently functionally linked to transcription regulation (Wang et al., 2023). H3K4me1 has been observed at active enhancers genome-wide together with H3K27ac (Creyghton et al., 2010). We find that H3K4me1, which is often associated with enhancers, is phosphorylated as efficiently as the unphosphorylated form of this peptide. Thus, methyl-phos switching could still operate at enhancers.

We did not observe changes in total levels of H3K9 and H3.1K27 methylation as rod-shaped chromosomes form. A previous study reported reduced chromosome size from *Suv39h dn* cells, while H3K27me was upregulated on these chromosomes (Djeghloul et al., 2023). However, our observations make it unlikely for heterochromatin to have a leading role in mitotic chromosome compaction.

Our analysis of phosphorylated peptides, where H3T3ph levels peak at approx. 30%, and genome-wide analysis by ChIP-seq reveal that Haspin phosphorylation of H3T3 appears not, as previously believed, to be an obligatory marker of centromeres (Kelly et al., 2010, Wang et al., 2010, Yamagishi et al., 2010, Harris et al., 2023) but instead is associated with heterochromatin. Our results reveal that methylation of K4 per se does not block Haspin phosphorylation of T3. Thus, the lack of phosphorylation of H3K4me2/me3 is likely because the mark is not in heterochromatin.

We were very surprised to note that although most centromeres were phosphorylated on T3 during mitosis, the centromeres of chromosomes Z and 5 were markedly devoid of this mark. We initially doubted our data and repeated the analysis using spike-in ChIP. The results obtained were virtually identical. Thus, it cannot be the case that centromere activity and targeting of the chromosomal passenger complex obligatorily requires phosphorylation of T3 by Haspin kinase (Kelly et al., 2010, Wang et al., 2010, Yamagishi et al., 2010, Jeyaprakash et al., 2011). Instead, Haspin is somehow marking heterochromatin in mitosis. Whether this serves a function – e.g. in targeting condensin or influencing the behavior of cohesin – remains to be determined. We note, however, that early studies of the localization of INCENP revealed that in early mitosis, the protein tended to cover the entire chromosomes before concentrating in the inner centromere during prometaphase and later (Cooke et al., 1987). The role of heterochromatin phosphorylation on T3, and whether the CPC has a role in early mitosis on chromosome arms remain subjects for future study.

One of the most surprising results of this study was that chromatin compaction and formation of rod-shaped chromosomes during prophase does not require core histone deacetylation. We had earlier shown that in the absence of condensin, rod-shaped mitotic chromosomes cannot form, but the total chromatin volume is not increased (Samejima et al., 2018). This suggested that something other than condensin must drive chromatin compaction. We then performed an in vitro study which revealed that core histones from mitotic cells assemble chromatin on a Widom array (Lowary and Widom, 1998) with more of a propensity to aggregate than that assembled from histones from interphase cells (Zhiteneva et al., 2017). Mass spectrometry analysis then went on to suggest that in addition to phosphorylation, a major difference between mitotic and interphase chromatin was deacetylation of core histones (Zhiteneva et al., 2017, Javasky et al., 2018). Others followed this study and argued that histone deacetylation is a critical factor driving chromosome compaction (Schneider et al., 2022).

Our present studies now strongly argue that those conclusions were incorrect. Indeed, measuring a number of different acetylated peptides, we find that rod-shaped chromosomes form at a time when histone deacetylation is barely detectible by mass spectrometry or immunoblotting. Importantly, a recent study from our lab revealed that in human cells, mitotic chromosomes reach the maximal concentration of nucleosomes (over 750 µM) by late prometaphase (Cisneros-Soberanis et al., 2024). This corresponds to the 30 minute time point in free-running mitotic entry experiments here.

Lacking the Cdk1^as^ system used here, previous studies had accumulated mitotic cells using inhibitors such as nocodazole or STLC (Zhiteneva et al., 2017, Javasky et al., 2018). Strikingly, we show here that such a mitotic delay results in a precipitous deacetylation of the histones. Our results reveal that at least as far as the initial stages of mitotic chromosome formation through prometaphase are concerned, bulk histone deacetylation is not required. Prolonged mitosis was previously shown to lead to an assembly of p53-containing complexes and cause p53 response in subsequent G_1_ (Meitinger et al., 2024), however the relationship between the cell cycle regulation and histone deacetylation remains to be determined.

Acetylation retained on mitotic chromatin might have a role in gene bookmarking (Behera et al., 2019), however genes constitute only a small percentage of the genome, so the levels of acetylation detected here are more likely to have another role in mitotic chromosome formation or segregation.

By far, the predominant histone modifications observed in mitosis are phosphorylations of H3T3, S10 and S28. Indeed, S10 phosphorylation is nearly quantitative. The methyl-phos switching observed between H3K9me3 and S10ph has been well documented, but this would be expected to occur only in heterochromatin and mitotic chromosomes still form if Aurora B kinase is inhibited (Adams et al., 2001, Ditchfield et al., 2003, Hauf et al., 2003). Furthermore, H3S10ph is installed in early prophase and persists throughout mitosis. We speculate that by decreasing the over- all positive charge of the chromatin, this modification might contribute to the bulk exodus of proteins from chromatin that occurs during mitotic progression (Samejima et al., 2022). What then of the delayed phosphorylation of S28 that occurs at around the time of nuclear envelope disassembly? It is tempting to speculate that an as-yet undiscovered methyl-phos switch involving H3K27 might be involved in releasing the condensed prophase chromosomes from their association with the nuclear envelope. It will be very interesting to screen amongst nuclear envelope proteins for any that can act as readers of methylated H3K27.

In addition to providing a baseline delineating the behavior of histone modifications in mitosis, the present study leaves us with no viable model for the nature of the process that dives the remarkable compaction of chromatin that occurs during mitosis. Clearly, this is a fertile area for future research.

## Limitations of the study

CDK1^as^ cells were accumulated at G_2_/M boundary with 1NM-PP1. Washout of 1NM- PP1 triggers rapid activation of CDK1 and mitotic entry, which may potentially affect the profile of histone modifications. Although histones are highly conserved between species, the exact modification profile may not be exactly the same between species and cell types.

Our LC-MS approach allowed for peptide profiling, corresponding to core histones H2A, H3 and H4. Trypsin digestion of propionylated H2B yields very long and highly charged peptides rendering impossible the unambiguous assignment of HCD fragment spectra. The same problem was encountered with two phospho-peptides: H2AT120ph and H3.3S28ph/S31ph (both of these are on the same peptide). Given the fact that H2B is the histone whose deacetylation was most dramatic in nocodazole-blocked cells (Zhiteneva et al., 2017), we validated our observations of deacetylation by western blotting of three of the five acetylation sites measured in that study. Lastly, our LC-MS measurements were performed without spike-in peptides, which makes the precise quantification difficult. We therefore performed spike-in ChIP-sequencing of H3Thr3ph, which is phosphorylated at 30% of all residues according to our LC-MS data.

## Author Contributions

NK designed experiments, performed cell culture and histone purification; MB designed and performed mass spectrometry analysis; MD designed and performed ChIP-seq experiments and immunofluorescence of H3T3ph; KS isolated all cell lines; IU and IS helped with experiments; IF helped with LC-MS analysis; NK, MB, MD, SW, IS, AI and WCE analysed data. LX and JRP synthesized 1NM-PP1. WCE and AI acquired resources and funding. NK, MD and WCE wrote the manuscript with contributions from all authors.

## Acknowledgments

We thank Hiroshi Kumira and Bryan Turner for the gift of antibodies; and Sam Corless and Alba Abad for helpful discussions and manuscript draft editing. We are grateful to Caitlin Reid for technical help and manuscript draft editing. We are grateful to Antoni Sieminski and the Centre for Statistics Drop-in Clinic at the University of Edinburgh for help with statistics. This work was funded by Wellcome grants 107022 and 221044 to WCE and 203149 to the Wellcome Centre for Cell Biology. This work was supported by funding for the Wellcome Discovery Research Platform for Hidden Cell Biology [226791] and we gratefully acknowledge support from the Bioinformatics core. AI lab was funded via grants from the Deutsche Forschungsgemeinschaft (DFG, 419067076 and 213249687). We are thankful to the Edinburgh Clinical Research Facility for the next-generation sequencing and Edinburgh Genome Foundry for assembling CENP-H construct used for establishing CENP-H cell line.

The authors declare no competing financial interests.

**Fig. S1. Purification of histones throughout short mitotic entry and nocodazole treatment.** A - A scheme of the short free-running mitotic entry experiment with one nocodazole treated sample. B – Images from the experiment with immunofluorescence against Lamin B1. Scale bar represents 5 μm. C – Quantification of mitotic phases from B. Bars represent mean ± standard deviation. n=5. D – Coomassie stained gel of purified histone proteins from the experiment.

**Fig. S2. Phosphorylation of S10, T3 and S28 on histone H3 in nocodazole block.** A, B and C – H3S10, H3T3 and H3.1S28 phosphorylation levels as the % of total peptide in the nocodazole experiment. Bars represent mean ± standard deviation. Individual replicates are shown with distinct colors. At G_2_ timepoint, an outlier replicate was excluded from the calculation of mean and standard deviation, as well as statistical analysis. n=5. For statistical comparison, Kruskal-Wallis test, followed by Conover-Iman test (in case of p < 0.05 in Kruskal-Wallis test) was performed. The p-values were adjusted with Benjamini-Hochberg procedure. D – A grouped barplot of the total mean peptide % of H3K9me1, me2, me3 and unme modifications in the nocodazole experiment. Error bars represent standard deviation. G – A grouped barplot of the mean % of H3K9me1, me2, me3 and unme which is phosphorylated (% of total phosphorylated peptides normalized to the total peptide % of these modifications). Error bars represent standard deviation. n=5. At G_2_ timepoint, an outlier replicate was excluded from the calculation of mean and standard deviation. E and H – The same as in D and G, but for H3K4me1, me2, me3 amd unme. At G_2_ timepoint, an outlier replicate was excluded from the calculation of mean and standard deviation of phosphorylation %. n=5. F and I – The same as in D/G and E/H, but for H3.1K27me1, me2, me3 and unme. At G_2_ timepoint, an outlier replicate was excluded from the calculation of mean and standard deviation of phosphorylation %, except H3.1K27unme S28ph%, where this replicate could not be considered an outlier.

**Fig. S3. H3K4me2 and H3K4me3 peptides are not phosphorylated on T3 during entry into mitosis.** A – Mean peptide % of H3K4me2 and me3 from the free-running mitotic entry experiment on a small scale (-0.25 – 2 %). n=4. Error bars represent standard deviation. For H3K4me2 % at 15 min, an outlier replicate was excluded from the calculation of mean and standard deviation. B – Mean % of H3K4me2 and me3 peptides, which is phosphorylated. n=4. Error bars represent standard deviation. C and D – The same as in A and B but for nocodazole experiment. No replicates were excluded from calculations. n=5.

**Fig. S4. Methylation enzymes and readers mostly dissociate from mitotic chromosomes, while phosphorylation enzymes do not.** – A, B, C– Chromatin enrichment of H3K9me, H3K4me and H3K27me writers from the free-running mitotic entry experiment of (Samejima et al, 2022). D, E, F and G, H, I – The same but for erasers and readers. J – The same for Aurora B kinase, which phosphorylates H3S10 and H3S28, and Haspin kinase (GSG2 in chicken), which phosphorylates H3T3. K – The same for H3T3, S10 and S28 phosphatases and their regulatory subunits.

**Fig. S5. Free-running mitotic entry experiment for ChIP-sequencing of key histone modifications.** Mitotic entry description for ChIP-sequencing. A – Images from the experiment with immunofluorescence against Lamin B1. Scale bar represents 5 μm. B – Quantification of mitotic phases from B. Bars represent mean ± standard deviation. n=4 (Two of the replicates include 2 technical replicates each). C – A Pearson correlation heatmap of differing replicates of H3T3ph ChIP-sequencing. Replicates 3 and 4 were performed with spike-in.

**Fig. S6. H3T3ph is broadly excluded from the TSS regions.** Heatmaps of normalized read densities of H3K4me1, H3K4me2, H3K4me3 and H3T3ph around the TSS regions (TSS ± 20 kb) throughout early mitosis.

**Fig. S7. H3T3ph is excluded from the centromere of non-repetitive chromosome Z.** A – A UCSC genome browser view showing the normalized ChIP-seq read densities of CENP-H (red), H3T3ph (dark blue), H3K9me3 (dark pink), H3K27me3 (dark orange) on Chr5 (upper panel). Regions shaded in pink highlight the centromeric region, which is shown in the lower panel at a higher magnification for a clearer view. B - Spike-in calibrated H3T3ph levels at the non-repetitive centromere of chromosome Z, as well as its p- and q-arms. Statistics was computed by fitting a linear mixed effect model of log(enrichment). “Timepoint” and “position along the chromosome“ were fit as fixed effects, and “sample” as a random effect. Gaussian distribution of models’ residuals was confirmed. Since T-distribution is symmetric, one-sided p value was calculated for the null hypothesis of no centromeric enrichment of H3T3ph. The hypothesis was not rejected (n.s.).

**Fig. S8. H3T3ph is enriched over the chromosome arms in addition to localizing to repetitive centromeres.** A – Snapshots of the UCSC genome browser view showing the normalized ChIP-seq read densities of H3K4me1 (light blue), H3K4me2 (dark green), H3K4me3 (violet), H3T3ph (dark blue), CENP-H (red), H3K9me3 (dark pink) and H3K27me3 (dark orange) on the whole Chr8 (Left) and Chr5 (Right). B – Immunofluorescence analysis of H3T3ph and ACA (anti-centromere antibodies) staining throughout mitotic entry. Scale bar represents 5 μm.

**Fig. S9. Nocodazole and MG132 treated mitotic entry allows capturing prometaphase cells at different lengths of mitosis.** A - A scheme of the nocodazole and MG132 treated mitotic entry experiment. B – Images from the experiment with immunofluorescence against Lamin B1. Scale bar represents 5 μm. C – Quantification of mitotic phases from B. Bars represent mean ± standard deviation. n=4. For the 4th replicate, immunofluorescence with Lamin B2 was performed. D – Coomassie stained gel of purified histone proteins from the experiment.

## Methods

### Cell culture, mitotic releases and cell filtering

All the cell lines used in this study are CDK1^as^ cells. The establishment of DT40 CDK1as cells and expression of OsTIR1 were described previously (Gibcus et al., 2018). Chicken DT40 CDK1^as^ cells (Gibcus et al., 2018) were grown in a log phase in RPMI media supplemented with 10% FBS and 5% chicken serum. Cells were split to 0.25 mln/ml 10-12 hours before 1NM-PP1 addition. 50 ml of cells per sample were blocked with 2 μM 1NM-PP1 for 13 hours.

G_2_ flask cells were filtered in a reusable filter unit (Nalgene) through 0.2 μm nitrocellulose filter and 1 ml of cells was crosslinked in the process of filtering for further immunofluorescence. The filter with cells was crumpled in 5 ml of 0.1 M H_2_SO_4_ and the 50 ml falcon containing it was placed on ice.

Cells for mitotic release were combined and washed 2 times with RPMI supplemented with 56 mM PIPES pH 7.0 (Samejima et al., 2022), after which they were resuspended in 2 ml RPMI-PIPES per sample and added to a flask with 48 ml RPMI-PIPES. The total cells washing and aliquoting time ranged from 6 minutes for the free-running mitosis and nocodazole experiments to between 6-7 minutes for nocodazole-MG132 treated mitosis. Cells were released into mitosis for denoted time in the incubator and collected in the last 30 seconds of the timepoint as described above for G_2_ cells. For nocodazole 120 minutes sample in nocodazole experiment, nocodazole (0.5 μg/ml) was added to the flask after 16 minutes release and cells were incubated until 120 minutes. For nocodazole and MG132 treated mitosis, nocodazole (0.5 μg/ml) or MG132 (20 μM) were pre-added to the flask before release.

### Cenp-H-Halo cell line establishment

Cenp-H-Halo cell line was established into the Cdk1^as^ background (Gibcus et al., 2018) using CRISPR/Cas9 technique to knock-in a Halo tag into the C-terminus of the gene. Knock-in constructs contain plasmids encoding a guide RNA and hCas9 were assembled by inserting double-stranded oligos (sense: CACCGCATTCAGAATCTCTTGTAGT and antisense: AAACACTACAAGAGATTCTGAATGC) into the pX330 which was a gift from F. Zhang (Addgene plasmid # 42230) (Cong et al., 2013) The 5’ and 3’ arms (500 bp each before/after stop codon) were synthesised by GeneArt. Then 5’ and 3’ arms, a Halo tag, as well as Hygromycin resistance cassette were assembled at Genome Foundry. 2 µg of knock-in construct and 6 µg of GuideRNA construct were transfected together into 4 x 10^6^ cells in the presence of 100 µl of buffer R. The cells were resuspended in 10 ml fresh medium and incubated. After 24 hrs, these cells were plated into 96 well plate with fresh medium supplemented with 1 µg/ml hygromycin. CENP-H is on the Z chromosome, thus, only one allele exists in DT40 cells. Integration of Halo tag was confirmed by imaging of cells with the Halo tag dye (JF549).

### Purifying histones

The cells were collected on nitrocellulose filters as described above. The filters containing the washed cells were placed in 50 ml falcons with 5 ml H_2_SO_4_ and rotated for 2 hours 30 minutes at 4°C for histone extraction. The supernatant was taken and the filter was discarded. The supernatant containing the histones was centrifuged 20000 x g 20 min, and the soluble fraction was collected in a 15 ml falcon. 0.1 M sulfuric acid solution with extracted histones was neutralized by the addition of 5 ml 1M Tris pH 8.0. Tris was slightly topped up (to 10 ml total volume), and NaCl (to 300 mM), NaEDTA pH 8.0 (to 2 mM), PMSF (to 0.25 mM) and DTT (to 1 mM) were added. SP Sepharose^TM^ Fast Flow beads (0.5 ml per sample) pre-equilibrated with loading buffer (50 mM Tris pH 8.0, 300 mM NaCl, 2 mM NaEDTA pH 8.0, 0.25 mM PMSF and 1 mM DTT) were mixed with approx. 11 ml of adjusted supernatant and rotated 1 hour at 4°C. The beads of each sample were washed 3 x 5 minutes with rotation at 4°C in 5 ml of wash buffer (50 mM Tris pH 8.0, 400 mM NaCl, 2 mM NaEDTA pH 8.0, 0.25 mM PMSF and 1 mM DTT) and elution was done for 5 minutes with rotation at 4°C in 1.5 ml elution buffer (50 mM Tris pH 8.0, 2 M NaCl, 2 mM NaEDTA pH 8.0, 0.25 mM PMSF and 1 mM DTT). The 1.5 ml eluate was collected and afterwards 90 μl 70% perchloric acid (4 % final concentration) was gradually added in 3 steps 30 μl each, with mixing in between. Histones were precipitated overnight, and the next day the samples were centrifuged 1 hour 15 minutes 18500 x g at 4°C. The pellets were washed 2x with 4% perchloric acid, 1x with 0.2% HCl in acetone and 1x with acetone. The pellets were dried and resuspended in 50 μl water supplemented with 0.25 mM PMSF. 10 μl were used for measuring the concentration in Bradford assay. 4x Laemmli loading buffer was added to the remaining histones to the final concentration of 1x and DTT was added to the final concentration of 0.1 M. These histones solutions were boiled and stored in -20°C before running on the gel.

### Running histone gels

The histone samples were boiled for 5 minutes at 95°C and resolved in a 18 % SDS-PAGE until the 6 kDa marker band reached the bottom of the gel. The gel was stained with Coomassie Imperial™ Protein Stain according to manufacturer’s instructions and destained in water overnight. The gel was imaged on Bio-Rad GelDoc Imaging System or Epson Perfection 3170 photo scanner.

### Immunofluorescence of Lamin during mitotic entry

1 ml of cells per sample were crosslinked with 1% formaldehyde for 10 minutes during filtering and quenched with 0.25 ml 2.5M glycine. Cells were washed in TBS and adhered to poly-L-lysine treated coverslips for 1 hour. Cells were crosslinked for 10 minutes at room temperature with 4% formaldehyde in PBS, prewarmed at 37°C, and permebealised with 0.15% Triton in PBS for 5 minutes. Cells were rinsed twice with PBS and blocked with 5% BSA in PBS for 30 minutes at room temperature. Cells were further rinsed in PBS and incubated with primary antibody in 5% BSA (AB16048 rabbit anti-LaminB1, batch 1022148-1 – 1:1000 dilution, or mouse anti-LaminB2, a gift from Erich Nigg - 1:1000 dilution). Cell were washed 3 times for 5 minutes with PBS, and incubated in the dark with secondary antibody (Goat anti-Rabbit IgG, Alexa Fluor™ Plus 488, A32731 or Goat anti-Mouse IgG (H+L) Alexa Fluor™ Plus 555, A32727, both 1:1000 dilution) in 5% BSA for 30 minutes. The cells were washed 2 x for 5 minutes with PBS, stained with Hoest 33342 2 μg/ml in PBS for 10 minutes, rinsed with PBS and mounted in ProLong™ Glass Antifade Mountant.

### Immunofluorescence of H3T3ph

DT40 CDK1^as^ cells were treated with 1NM-PP1 for 13 hours, and then cells were released by washing out the 1NM-PP1 with fresh medium. Cells were fixed with 1% formaldehyde for 10 minutes at room temperature. Fixed cells were then quenched with freshly prepared 1.25 M glycine for 5 minutes and washed three times with ice-cold PBS and adhered to poly-L-lysine coated coverslips. Cells were permeabilized with 0.2% Triton-X100 for 5 min. After washing with PBS, the cells were incubated for 1 hour with 5% BSA in PBS and then with primary antibodies, H3T3ph (16B2; mouse; diluted 1:500 in blocking buffer) and ACA (human; Antibodies Incorporated; 15-234; diluted 1:500 in blocking buffer), in 1% BSA and 0.1% Tween-20 in PBS overnight at 4°C. After washing twice with 0.1% Tween-20 in PBS, cells were incubated for 1 h with fluorophore-conjugated secondary antibodies: anti-mouse Alexa Fluor^®^488 (Invitrogen A32723) and anti-human Alexa Fluor^®^647 (Invitrogen A21445). Cells were rinsed three times in 0.1% Tween-20 before DNA staining and mounting with DAPI in Vectashield Plus (Vector Laboratories).

### Microscopy

Slides with Lamin staining were imaged on a DeltaVision microscope using SoftWorx 6.0 software (Applied Precision/GE Healthcare) with Olympus UPlanXApo 100X oil objective, NA 1.45. Images were acquired as z-stacks at 0.5 μm intervals, and single sections were shown in the figure panels. Exposures for images were adjusted individually. Images were processed in ImageJ. For mitosis stage classifications, slides were viewed in DAPI and FITC/TRIC channels under the microscope. Cells with intact nuclear lamina were classified as interphase or prophase, but in case of the break in the nuclear envelope the cell was staged as prometaphase. Images were not deconvolved, as if breaks in prophase lamina appeared upon deconvolution, one would not be able to exclude artifacts.

For H3T3ph staining, images were acquired as z-stacks at 0.2 μm intervals and deconvolved using SoftWoRx. Maximum intensity projections were shown. Exposures for images were adjusted individually.

### Western blotting

Histones were resolved on 18% SDS-PAGE, as described above. The transfer to the nitrocellulose membrane (Amersham) was performed at 4°C for 2 h at 100 V. The membrane was rinsed with 0.2% Tween in PBS (PBST) and blocked in 5% milk in PBS for 1 hour at room temperature. The membranes where incubated with primary antibody (rabbit anti-H2B5ac, 12799T, Cell Signalling, 1:1000; rabbit anti-H2BK12K15ac, R209, from Bryan Turner, 1:400; rabbit anti-H4K8ac, R403, from Bryan Turner, 1:400; rabbit anti-H2B, ab1790, Abcam, 0.2 μg/ml; rabbit anti-H4, ab10158, Abcam, 1 μg/ml) in PBST with 1% milk overnight at 4°C. Next day the membrane was washed twice with PBST and incubated with secondary antibody (anti-rabbit Alexa 800, A32735, Invitrogen, 1:10000 or 926-32213 IRDye 800CW anti-rabbit IgG, LICORbio™, 1:10000) in PBST with 1% milk for 1 h at room temperature. The membrane was further washed twice with PBST and imaged on the LiCor Odyssey CLx Infrared Imaging System (LICORbioTM). Images were quantified and processed with Image Studio™ Software (LICORbioTM). For quantification, a rectangle was drawn around the band and intensity of rectangles was normalized against G2, as well as loading control for respective lane.

### Histones preparation for mass spectrometry

Histone analysis by mass spectrometry was performed as described previously (Volker-Albert et al., 2018). 1 μg of histones were resuspended in an appropriate volume of 1x Laemmli buffer to reach a final volume of 10 μL and boiled at 95°C for 5min. Proteins were resolved on a precast 4-20% polyacrylamide gels (SERVAGel™ TG PRiME™) which were then stained with InstantBlue Coomassie. Histones bands were excised (between 11-17 kDa), washed once with water and de-stained three times by adding 50 μL 50 mM NH_4_HCO_3_ and 50 μL of acetonitrile and incubating at 37°C for 30 min. Gel pieces were washed twice with water before being dehydrated three times with 150 μL of acetonitrile. 5 μL of propionic anhydride were used for in gel histone acylation followed by the addition of 10 μL of 100 mM NH_4_HCO_3_ and 35 μL of 1M NH_4_HCO_3_ and incubation at 37°C for 45 min. Subsequently, samples were washed five times with 150 μL of 100 mM NH_4_HCO_3_, 150 μL of water and 150 μL of acetonitrile. Gel pieces were re-hydrated with 20 μL of trypsin solution (25 ng/μL trypsin (Promega) in 100 mM NH_4_HCO_3_) and incubated at 4°C for 30 min. After the addition of 50 μL of 50 mM NH_4_HCO_3_, in gel digestion was achieved by overnight incubation in the thermomixer at 37°C at 550 rpm. Digested peptides were extracted by the addition of 50 μL of 50% acetonitrile 0.25% TFA (twice) and 50 μL of 100% acetonitrile (twice) incubating 10 min at RT. The extracted peptides were filtered using a C8 Empore disk stagetips and the obtained flow through evaporated to dryness using a SpeedVac concentrator Plus (Eppendorf). Dried peptides were resuspended in 15 μL of 2% ACN 0.3% TFA and stored at -20°C until mass spectrometry analysis. Pooled quality control samples were generated by combining 1 μL of each reconstituted sample and analyzed repeatedly at the beginning and within the sequence.

### Instrumentation of LC-MS

1 μL was injected in an Ultimate 3000 RSLCnano system (ThermoFisher Scientific) and separated in a 25 cm analytical column (75 µm ID, 1.6 µm C18, Aurora-IonOpticks) with a 90 min gradient from 3 to 43% ACN in 0.1% formic acid at a flowrate of 300 nl/min before being electrosprayed into QExactive HF mass spectrometer (ThermoFisher Scientific). Survey full-scan MS spectra (from m/z 250 to 900) were acquired with resolution R = 60,000 at m/z 400 (AGC target of 3e6). Top 10 most intense ions (charge state between 2 and 5) were sequentially isolated with an isolation window of 2.0 m/z to a target value of 2e5 and fragmented at 27% normalized collision energy and acquired with resolution R= 15,000 at m/z 400 (max IT 60 ms). Ion spray voltage was set to +1.5 kV with no sheath and auxiliary gas flow and heated capillary temperature was set to 250°C.

### Data analysis

Raw data from mass spectrometry was analyzed using Skyline software (MacCoss Lab, University of Washington, Seattle, WA, USA) (Pino et al., 2020). Automatic selection of doubly and triply charged peptides masses was manually curated based on the relative retention times and fragmentation spectra with results from Proteome Discoverer 1.4 software (ThermoFisher Scientific). Integrated peak areas were exported and used to calculate relative levels of each PTMs using R environment (R Core Team, 2022), based on the formula given in (Volker-Albert et al., 2018). The resulting data was analyzed and plotted in R.

### ChIP-sequencing

For chromatin immunoprecipitation, each sample was prepared from 10 million Chicken DT40 CDK1^as^ cells for each condition. The H3K4me1 (CMA302), H3K4me2 (CMA303), H3K4me3 (CMA304) and H3T3ph (16B2) antibodies were kindly provided by Prof. Hiroshi Kimura. The antibody specificity was analysed (Kimura et al., 2008). Dynabeads™ M-280 Sheep Anti-Mouse IgG (Invitrogen: 11201D; 40 μl for each sample) for H3K4me1, H3K4me2, H3K4me3, H3T3ph and Dynabeads™ Protein A for Immunoprecipitation (Invitrogen: cat no. 10002D; 25 μl for each sample) for H3K9me3 (abcam, ab8898) were used to perform the ChIP experiment. For CENP-H ChIP analysis we used DT40 CDK1^as^-CENP-H-Halo cells. CENP-H was immunoprecipitated against the Halo protein, where 4 million *Drosophila melanogaster* S2-CID-Halo cells (Sacristan et al., 2024) were used as a spike-in control. In H3T3ph ChIP analysis, we used 0.3 million RPE-CDK1^as^ cells (Cisneros-Soberanis et al., 2024) as a spike-in control.

Briefly, DT40 CDK1^as^ cells were treated with 1NM-PP1 for 13 hours and then cells were released by washing out the 1NM-PP1 with fresh medium. DT40 CDK1^as^, S2-CID-Halo, and RPE CDK1^as^ cells were fixed separately with 1% formaldehyde for 10 minutes at room temperature under shaking condition. Fixed cells were then quenched with freshly prepared 1.25 M glycine for 5 minutes at room temperature on a shaker and washed with ice-cold PBS three times. The cells were then snap-frozen using dry ice and ethanol and preserved at -80°C for ChIP-seq.

For antibody binding, Dynabeads were washed with PBS supplemented with 1% BSA, and incubated with anti-histone antibodies for 4 hours at 4 °C. Cell pellets were thawed on ice and mixed at a specific ratio. The cells were then lysed with 120 μl of cold lysis buffer (1% SDS, 10 mM EDTA, 50 mM Tris-HCl (pH 8.1), 0.1 mM DTT, and protease inhibitor cocktail (Sigma)) by incubating for 16 minutes on ice. Afterwards, 400 µl of IP dilution buffer (1% Triton X-100, 2 mM EDTA, 20 mM Tris-HCl (pH 8.1), 150 mM NaCl, 0.1 mM DTT, and protease inhibitor cocktail) was added. To shear the genomic DNA, cell lysates were then sonicated for 16 cycles (30 sec on/off) at maximum power using Bioruptor® Plus sonication device. After centrifugation to remove debris, Triton X-100 was added to the lysates to achieve a final concentration of 1% for IP. 10% of the sheared chromatin was set aside for Input, and the rest was incubated with specific antibodies or equilibrated Halo-Trap Agarose beads (ChromoTek) for overnight at 4°C on a rotating wheel. The beads were washed at 4°C with three different wash buffers: wash buffer 1 (1% Triton X-100, 2 mM EDTA, 20 mM Tris-HCl (pH 8.1), 150 mM NaCl, 0.1 mM DTT and protease inhibitor cocktail), wash buffer 2 (50 mM Hepes pH 7.9, 1% Triton X-100, 1 mM EDTA, 50 mM Tris-HCl (pH 8.1), 500 mM NaCl, 0.1% Na-deoxycholate, 0.1% SDS, 0.1 mM DTT and protease inhibitor cocktail), wash buffer 3 (20 mM Tris-HCl pH 8.0, 1 mM EDTA, 250 mM LiCl, 1% NP-40, 0.1% Na-deoxycholate, 0.1 mM DTT and protease inhibitor cocktail). Finally, beads were washed with TE buffer (1 mM EDTA, 10 mM Tris-HCl pH 8.0) twice, and chromatin was extracted by incubating the beads with extraction buffer (0.1 M NaHCO_3_, 1% SDS) twice at 65°C on a thermomixer for 15 mins. For reverse crosslinking, IP-DNA and Input-DNA were incubated with 200 mM NaCl at 65°C overnight. Then, they were treated with RNase-A at 37°C for 1 hour and then with proteinase K at 65°C overnight. DNA was purified by using the QIAquick PCR Purification Kit. After quantification with the Qubit assay, ChIP-seq libraries were prepared using the NEBNext® Ultra™ II DNA Library Prep Kit for Illumina® with NEBNext® Multiplex Oligos for Illumina® (Index Primers Set 1) according to the manufacturer’s protocol. The library size distribution was analysed by Agilent Bioanalyzer with High Sensitivity DNA chip and submitted for Illumina deep sequencing (100 bp pair-end) at the Edinburgh Clinical Research Facility. Sequencing was performed on the NextSeq 2000 platform using the NextSeq 1000/2000 P2 Reagents (100 cycles).

## ChIP-sequencing analysis

### Genome Reference and Annotation

All read alignments and downstream analyses were performed against the *Gallus gallus* (chicken) telomere-to-telomere (T2T) reference genome assembly GGswu, accession GCA_024206055.2, downloaded from NCBI https://www.ncbi.nlm.nih.gov/datasets/genome/GCA_024206055.2/. This assembly provides a complete gapless representation of all chromosomes (Huang et al., 2023).

### Analysis of ChIP-seq data

Data processing was performed using a reproducible ChIP-seq workflow implemented with Snakemake (v6.15.1) to automate and document the analysis from raw sequencing data through peak calling and genome visualisation.

ChIP-seq libraries were sequenced using paired-end Illumina chemistry with a read length of 2 × 51 bp. Adapters were trimmed using Cutadapt v1.18 (Martin, 2011) with -- minimum-length 20 and --nextseq-trim=20. Quality of raw and trimmed reads was assessed using FastQC v0.11.9 (Andrews, 2010) and summary reports were aggregated using MultiQC v1.26 (Ewels et al., 2016).

Trimmed reads were aligned to the GGswu genome using BWA-MEM v0.7.16 (Li and Durbin, 2009) with the -M flag to mark shorter split hits as secondary. The resulting SAM files were filtered, sorted and indexed using Samtools v1.17 (Danecek et al., 2021), and alignment statistics were collected via samtools flagstat. Reads mapped to the combined GGswu_hg38 reference were separated by genome. Alignments were subjected to the following filtering steps: Removal of secondary (0x100), supplementary (0x800), and unmapped (0x4) reads (-F 2304), retention of properly paired reads (-f 3).

Signal tracks were generated using deepTools v3.5.0 (Ramirez et al., 2016). bamCoverage was used to compute normalised coverage in bins of 1 bp using the BPM (bins per million reads) method. bamCompare was used to compute log_2_ enrichment profiles for IP over input control, using exact scaling and BPM normalisation.

Further tracks were generated to represent the occupancy ratio (OR) of the IP for experiments containing spike-in. The OR was calculated using the protocol outlined in (Hu et al., 2015).

The ChIP-seq dataset was further analysed with Galaxy (usegalaxy.org) (The Galaxy Community, 2024) and R (R Core Team, 2022). For the ChIP-seq heatmap and enrichment plotting, the signal was quantified over defined windows around ± 20 kb from the TSS using deepTools computematrix (Version 3.5.4) by using the following parameters: 50 bp non-overlapping bins used for averaging the score over the length of the regions (--binSize), -- sortRegions was in descending order, --sortUsing was mean, --missingDataAsZero was 0 and --skipZeros was yes. The output was a matrix of signal values per region per sample, which was plotted via the plotHeatmap tool (Version 3.5.4) by using the following parameters: -- sortRegions in a descending order, --sortUsing was mean and clustering was performed via k-means algorithm (one cluster). Profile plots of each heatmap were calculated as the values of the columns in the matrices generated by deepTools and plotted as a group for H3K4me1, H3K4me2, H3K4me3 and H3T3ph where each group has a 5 timepoints dataset (G_2_, 5 min, 7.5 min, 15 min, 30 min) by using --perGroup as yes.

The average value of bigwig data from two replicates was calculated with the bigwigAverage tool (Version 3.5.4). These datasets were used to calculate the genome-wide enrichment signals by using the multiBigwigSummary tool (Version 3.5.4) with parameters 10000 bp bin size (--binSize) and --distanceBetweenBins as 0. The correlation matrix was prepared with the plotcorrelation tool with the following parameters: --corMethod was Pearson, --removeOutliers was no and --skipZeros was yes (Ramirez et al., 2016). A correlation heatmap was plotted via R.

Based on CENP-H map, Centromere, p-Arm and q-Arm were determined. By using the multiBigwigSummary tool (Version 3.5.4), H3T3ph enrichment was calculated in specific regions with parameters 10000 bp bin size and --distanceBetweenBins as 0. The values were plotted in R.

## Data availability

Histone mass spectrometry data are deposited to the ProteomeXchange Consortium via the PRIDE20 partner repository with the accession number PXD066101. ChIP-sequencing data are deposited to the Gene Expression Omnibus database with the accession number GSE302647.

